# Variation in relaxation of non-photochemical quenching in a soybean nested association mapping panel as a potential source for breeding improved photosynthesis

**DOI:** 10.1101/2020.07.29.201210

**Authors:** Steven J. Burgess, Elsa de Becker, Stephanie Cullum, Isla Causon, Iulia Floristeanu, Kher Xing Chan, Caitlin E. Moore, Brian W. Diers, Stephen P. Long

## Abstract

Improving the efficiency of crop photosynthesis has the potential to increase yields. Genetic manipulation showed photosynthesis can be improved in Tobacco by speeding up relaxation of photoprotective mechanisms, known as non-photochemical quenching (NPQ), during high to low light transitions. However, it is unclear if natural variation in NPQ relaxation can be exploited in crop breeding programs. To address this issue, we measured NPQ relaxation in the 41 parents of a soybean NAM population in field experiments in Illinois during 2018 and 2019. There was significant variation in amount and rate of fast, energy dependent quenching (qE) between genotypes. However, strong environmental effects led to a lack of correlation between values measured over the two growing season, and low broad-sense heritability estimates (< 0.3). These data suggest that either improvements in screening techniques, or transgenic manipulation, will be required to unlock the potential for improving the efficiency of NPQ relaxation in soybean.

**Table of Abbreviations:** 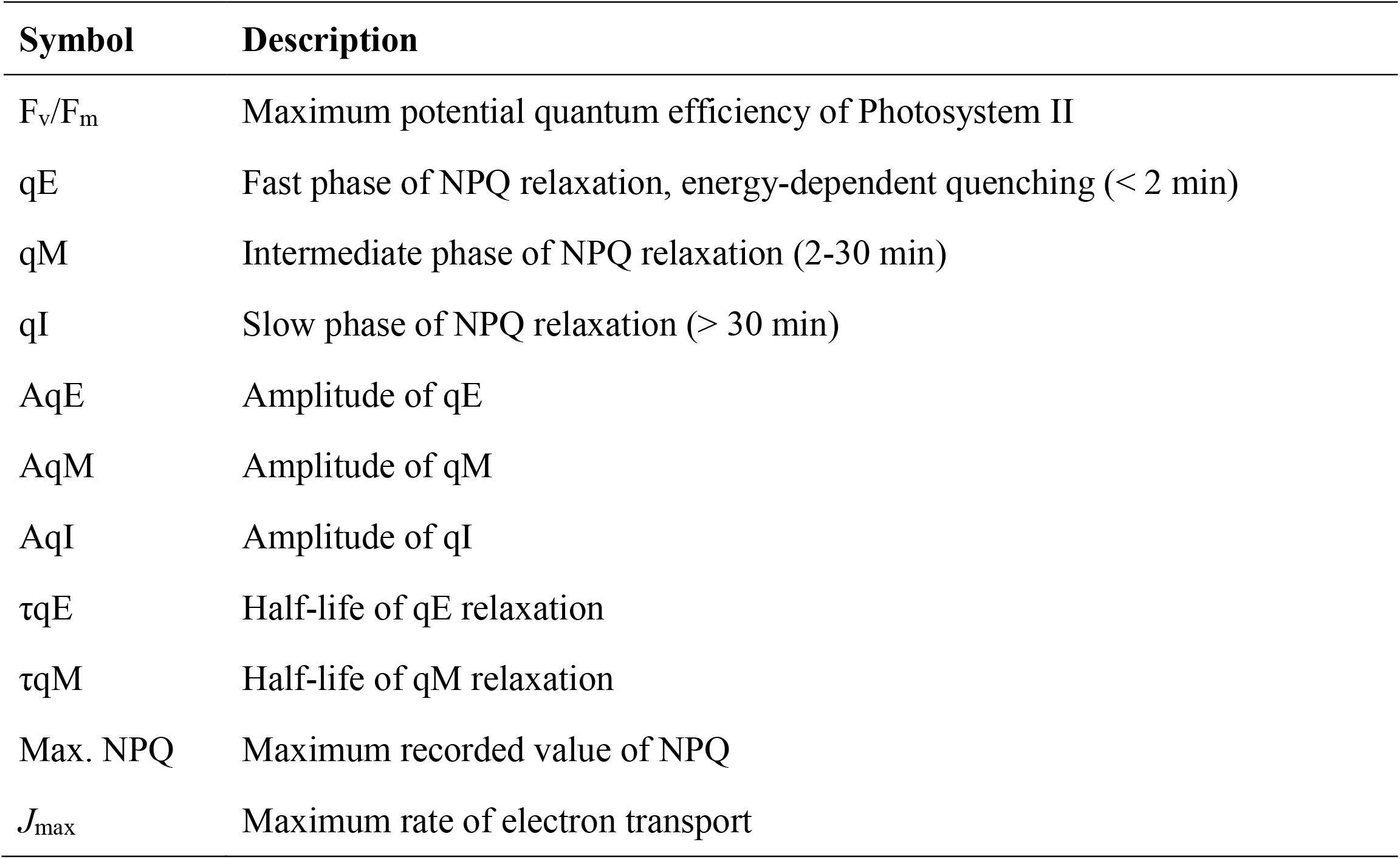

## Introduction

Soybean is a major source of vegetable protein worldwide. Improved agronomic practices and intensive breeding efforts led to sustained yield increases of 23.3 kg ha^−1^ y^−1^ between 1924 and 2012 in the USA (Specht et al., 2014). However, further genetic improvements are required to meet future demands without expanding the area of land under cultivation and causing environmental degradation (Grassini et al., 2013).

The maximum achievable yield in the absence of abiotic or biotic stress is known as the yield potential, and can be defined as:

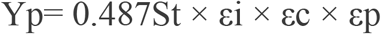

Where St is the total solar incident radiation, 48.7 % of which is photosynthetically active (PAR); canopy light interception (ε_i_) is the amount of solar radiation captured over a growing season; solar conversion efficiency (ε_c_) is the ability to convert captured solar energy into seed biomass; and harvest index (ε_p_) is the proportion of above-ground biomass that is partitioned into seed (Zhu et al., 2010).

Research suggests ε_p_ (Morrison et al., 1999; Koester et al., 2016) and ε_i_ (Dermody et al., 2008; Koester et al., 2014) are close to their theoretical maximum in modern soybean genotypes. In contrast, ε_c_ is calculated to be between 2-4 % (Koester et al., 2014), which is less than half the theoretical maximum (Zhu et al., 2010) making it an attractive target for improvement. Several strategies have been proposed to increase ε_c_ (Zhu et al., 2010; Long et al., 2019) and experiments in *Nicotiana tabaccum* have shown it is possible by genetic modification (Lefebvre et al., 2005; Kromdijk et al., 2016; South et al., 2019).

One strategy to improve ε_c_ focuses on altering photoprotection (Kromdijk et al., 2016). Photodamage can occur when the amount of light absorbed exceeds the rate at which it is used for the fixation of carbon, also known as photochemical quenching. Excess absorbed light energy can be dissipated by non-photochemical quenching (NPQ), reducing the likelihood that damaging reactive oxygen species are formed (Murchie and Lawson, 2013; Ruban, 2016). In a field environment, leaves within a canopy are exposed to instantaneous fluctuations in light caused by intermittent cloud cover and wind induced leaf movements. This requires rapid activation and deactivation of NPQ to adjust to altering light levels and reduce field recombination-induced photodamage (FRIP) caused by ^1^O_2_ production (Davis et al., 2016). However, a delay in relaxing NPQ during high to low light transitions is predicted to result in unnecessary dissipation of energy, reducing the efficiency of photosynthesis (Zhu et al., 2004). To address this, Kromdijk et al. (2016) manipulated the expression of three genes involved in photoprotection, including Zeaxanthin epoxidase (ZEP), Violaxanthin de-epoxidase (VDE) and Photosystem II subunit S (PsbS), showing it is possible to speed up the rate of activation and relaxation of NPQ, leading to improved efficiency of photosynthesis and increased biomass accumulation (Kromdijk et al., 2016).

NPQ is a collection of processes, including fast, energy dependent quenching (qE)(Krause et al., 1982), medium term processes (qM): zeaxanthin dependent quenching (qZ)(Nilkens et al., 2010) and state transitions (qT)(Quick and Stitt, 1989), and long term quenching: photoinhibition (qI)(Krause, 1988) and photoinhibition independent quenching (qH)(Malnoë et al., 2018).

However, it is unclear what is an ideal amount of photoprotection, measured as the amplitude of qE and qM, and if it can be optimized, as it is likely to vary between species and environment. Too much photoprotection will unnecessarily dissipate light energy that could be used for photosynthesis, while too little will result in photodamage, reducing photosynthetic efficiency. Therefore the best genotypes will have an empirically determined optimum level of NPQ, relax photoprotection quickly and experience minimal photodamage, thereby maximizing photosynthetic efficiency.

From a breeding perspective, it is important to assess both the degree of variation available in germplasm and the extent to which this variation is heritable in order to determine whether a trait can be improved through breeding. Previous studies on diverse genotypes in rice (Kasajima et al., 2011; Wang et al., 2017) and arabidopsis (Jung and Niyogi, 2009; Rungrat et al., 2019) have demonstrated the existence of substantial variation in NPQ within species, but focused only on maximum NPQ (Wang et al., 2017) or qE (Jung and Niyogi, 2009; Kasajima et al., 2011; Rungrat et al., 2019) and did not assess heritability.

Chlorophyll fluorescence was evaluated in two studies with field grown soybean, using canopy reflectance to calculate photochemical reflectance index (PRI), which is a proxy for NPQ (Herritt et al., 2016), and OJIP transients to look at variation in fluorescence kinetics (Herritt et al., 2018). Estimates of the broad sense heritability of traits (*H*^2^), ranged from 69 % for PRI and 36.8 % for dark fluorescence values F0, to 0.4 % for energy flux for electron transport per reaction center. However, the individual components of NPQ relaxation was not assessed by Herritt *et al.* (2016, 2018). Therefore, the extent to which individual components of NPQ co-vary in natural populations and whether they can be easily selected in breeding programs remains unclear.

Here, rates of NPQ relaxation in the 41 parents of the soybean nested association mapping (NAM) population (Song et al., 2017; Diers et al., 2018) were measured and a double exponential curve was fit (Dall’Osto et al., 2014) to calculate the amplitude of qE (AqE), qM (AqM), qI (AqI), and the half-life of qE (τqE) and qM (τqM) relaxation. This population includes 18 elite breeding genotypes, 15 diverse breeding genotypes that contain exotic diversity, and eight exotically sourced plant introductions. These data were used to test the hypothesis that (1) there is significant diversity in rates of NPQ relaxation between soybean genotypes and (2) individual components of NPQ relaxation vary independently between parents. The heritability of NPQ relaxation components were then calculated to assess the potential for breeding improved solar conversion efficiency.

## Methods

### Plants and growth conditions

The 41 parents of the Soybean Nested Association Mapping (NAM) population (Song et al., 2017, Diers et al., 2018) were grown in the field at the Crop Sciences Research and Education Center at the University of Illinois at Urbana-Champaign in 2018 and 2019. Seeds were planted on 23 May 2018 and 6 June 2019 in 1.2 meter long single-row plots with a 0.75 m row spacing at a rate of 40 seed m^−1^. The experiment was arranged in a randomized complete block design with five replicates and standard agronomic practices were employed.

### Meteorological data collection

Meteorological variables were measured every 30-minutes by a weather station at the University of Illinois Energy Farm, approximately 1 km from the Crop Sciences Research and Education Center (latitude 40.062832, longitude −88.198417). Air temperature (Ta, °C) and relative humidity (RH, %) were recorded by a HMP45C probe (Campbell Scientific, Logan, UT, USA), and incoming shortwave radiation (Fsd, W m^−2^) was from a CNR1 radiometer (Kipp & Zonen, The Netherlands), both instruments were installed 4 m above the ground. Rainfall (mm) was obtained from the Illinois Water Survey at 30-minute time step (Illinois State Water Survey, 2020). Ta and RH were used to calculate saturation vapor pressure (es) and actual vapor pressure (ea) for each 30-minute period, which were then used to calculate vapor pressure deficit (VPD, kPa) as per Equations 1-3:

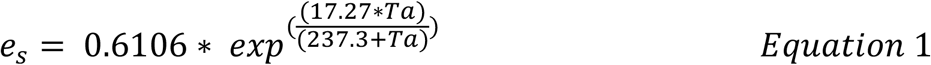

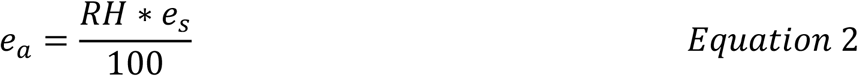

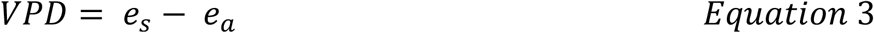

Occasional gaps in meteorological data are inevitable when measuring at these time scales, so data gaps were filled where needed following (Isaac et al., 2017), whereby an artificial neural network was used to generate a complete time-series with external data sourced from the University of Illinois Willard Airport weather station (station ID: 725315-94870) and ERA-interim data from the European Centre for Medium Range Forecasts (Dee et al., 2011). In total, less than 5 % of data required gap filling. Daily summary values were then calculated for each variable, with Ta and VPD presented as daily mean, Fsd as daytime-only (i.e. 06:00-18:00) mean, and rainfall as a daily sum. A 7-day rolling mean was calculated to aid with data visualization.

### Chlorophyll fluorescence analysis

Plants were sampled on the 19th of July 2018 and the 26^th^ July 2019, when at the R1-R3 developmental stage. Three 4.8 mm leaf disks were collected from the upper-most mature leaf of each replicate strip in the field using a cork borer (Humboldt H9663; Fisher Scientific 07-865-10B).

In 2018 leaf disks were placed face down in clear 96 well plates and held in position with a moist sponge cut to the size of a well. Plates were transported back to the laboratory for overnight dark incubation to allow for relaxation of long term NPQ. In 2019 leaf disks sampled in the field were floated on dH_2_O in a 24 well plate which was sealed with parafilm for transportation back to the laboratory for ease of manipulation. Leaf disks were then transferred to a square petri dish lined with wet filter paper, and plates sealed with parafilm and wrapped in aluminum foil for overnight incubation.

Measurements were taken with modulated chlorophyll fluorescence imaging systems, in 2018 using a CF imager (Technologica, UK) and in 2019 a FluorCam FC 800-C (PSI Systems, Czech Republic in 2019). Briefly, overnight incubated disks were subjected to 10 min illumination at 1000 μmol m^−2^ s^−1^ white light (6500 K) followed by 50 min of darkness. *F* m, was determined by applying saturating pulses (4000 μmol m^−2^ s^−1^ white light) at 9, 40, 60, 80, 100, 120, 160, 200, 240, 300, 360, 420, 480, 540 and 598 s after the actinic light was turned on, and at 1, 2, 4, 6, 8, 10, 14, 18, 22, 26, 32, 38, 44 and 50 min after the actinic light was turned off. The background was excluded automatically and NPQ values at each pulse were calculated.

In 2018, NPQ values were calculated for each time point using FluorImager v2.305 with custom MatLab R0218b scripts (see accession numbers) and in 2019 using the software FluorCam7 v.1.2.5.3. NPQ relaxation parameters AqE, AqM, AqI, τqE and τqM were then calculated by fitting a double exponential function to measured NPQ values following shut off of the actinic light, according to Equation 4 (Dall’Osto et al., 2014):

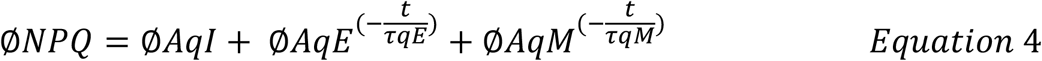

using the fit function in MatLab R0218b (see accession numbers), where t is the measured fluorescence at a given timepoint. Maximum NPQ values are defined as the maximum value reached during the 10 min illumination.

### Statistical Analysis

After visual inspection of data distributions, outliers were defined and removed if values were outside 1.5 x the interquartile range (IQR). Statistical significance was determined by one-way ANOVA with a post-hoc Games-Howell test at an alpha level 0.05 using the R package ‘userfriendlyscience 0.7.2’ (Peters, 2018) and custom R-scripts (see accession numbers). Pearson correlations between NPQ relaxation parameters AqE, AqM, AqI, τqE and τqM, and between years, were determined using the R package ‘PerformanceAnalytics 2.0.4’ (Peterson et al., 2020).

### Broad sense heritability calculation

The mean of technical replicates taken from genotypes in each year was used to estimate the heritability of NPQ relaxation parameters. Variance components were determined using a linear mixed effects model (lmer) function in the R package ‘lme4 1.1-23’ (Bates et al., 2015), with genotype as a random effect and environmental variance estimated from the residuals. Broad-sense heritability was then calculated according to Equation 5:

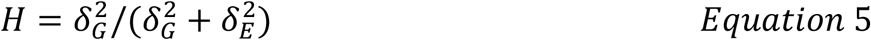

Where *H^2^* is broad sense heritability, δ^2^_G_ is genotypic variance and δ^2^_E_ is environmental variance.

### Accession numbers

Raw chlorophyll fluorescence imager files and scripts are deposited on figshare and can be accessed at https://doi.org/10.6084/m9.figshare.c.5075717.

## Supplemental Data Files

Supplemental Data 1: NPQ relaxation parameters calculated for the 41 parents of the Soybean Nested Association Mapping panel

Supplemental Data 2: ANOVA and Games-Howell posthoc test results comparing NPQ relaxation kinetics between 41 founder lines of Soybean NAM population

Supplemental Figure 1: NPQ relaxation kinetics of the 41 parents of the soybean NAM population in 2018.

Supplemental Figure 2: NPQ relaxation kinetics of the 41 parents of the soybean NAM population in 2018.

Supplemental Figure 3: Pearson correlation between NPQ parameters of the 41 parents of the soybean NAM population in 2018.

Supplemental Figure 4: Pearson correlation between NPQ parameters of the 41 parents of the soybean NAM population in 2019.

Supplemental Figure 5: Pearson correlation between measured AqE values of 41 parents of the soybean NAM population in 2018 and 2019.

Supplemental Figure 6: Pearson correlation between measured AqM values of 41 parents of the soybean NAM population in 2018 and 2019.

Supplemental Figure 7: Pearson correlation between measured AqI values of 41 parents of the soybean NAM population in 2018 and 2019.

Supplemental Figure 8: Pearson correlation between measured τqE values of 41 parents of the soybean NAM population in 2018 and 2019.

Supplemental Figure 9: Pearson correlation between measured τqM values of 41 parents of the soybean NAM population in 2018 and 2019.

Supplemental Figure 10: Pearson correlation between measured maximum NPQ values of 41 parents of the soybean NAM population in 2018 and 2019.

## Results

NPQ relaxation was measured for the 41 parents of the soybean NAM population over two growing seasons in Urbana, IL, USA (2018 and 2019), to investigate genotypic and environmental variation. Trends between genotypes were compared between years, rather than measured values, as different chlorophyll fluorescence imaging systems were used (see methods). In both years, the amplitude of NPQ attributable to fast, medium and slow relaxing phases was determined to be qE > qM > qI for all parents (Figure 1 and 2; Supplemental Figure 1 and 2, Supplemental Data 1).

**Figure 1:**
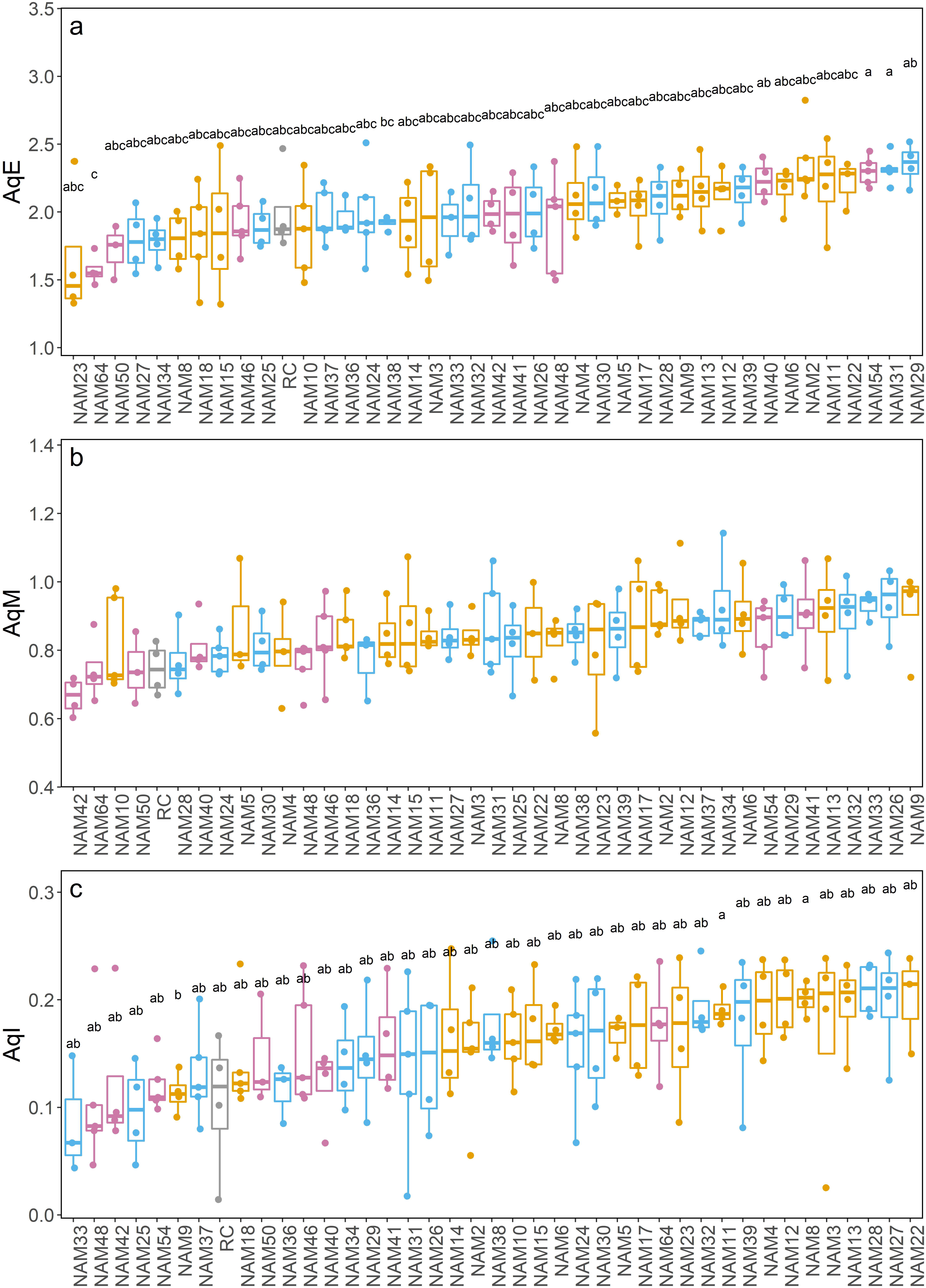
NPQ relaxation kinetics of the 41 parents of the soybean NAM population in 2019. Box plots representing (a) AqE, (b) AqM, (c) AqI, displaying mean and standard deviation. Data points for the mean of n=3-5 (See Supplemental Data 1 for details) biological replicates are displayed. Colors correspond to genotype groups, including: Reference genotype (grey), EL = elite genotype (orange), BX = breeding genotype with exotic ancestry (blue), PI = plant introductions (magenta). Letters denote significant groupings (α=0.05) based on Games-Howell posthoc test.

**Figure 2:**
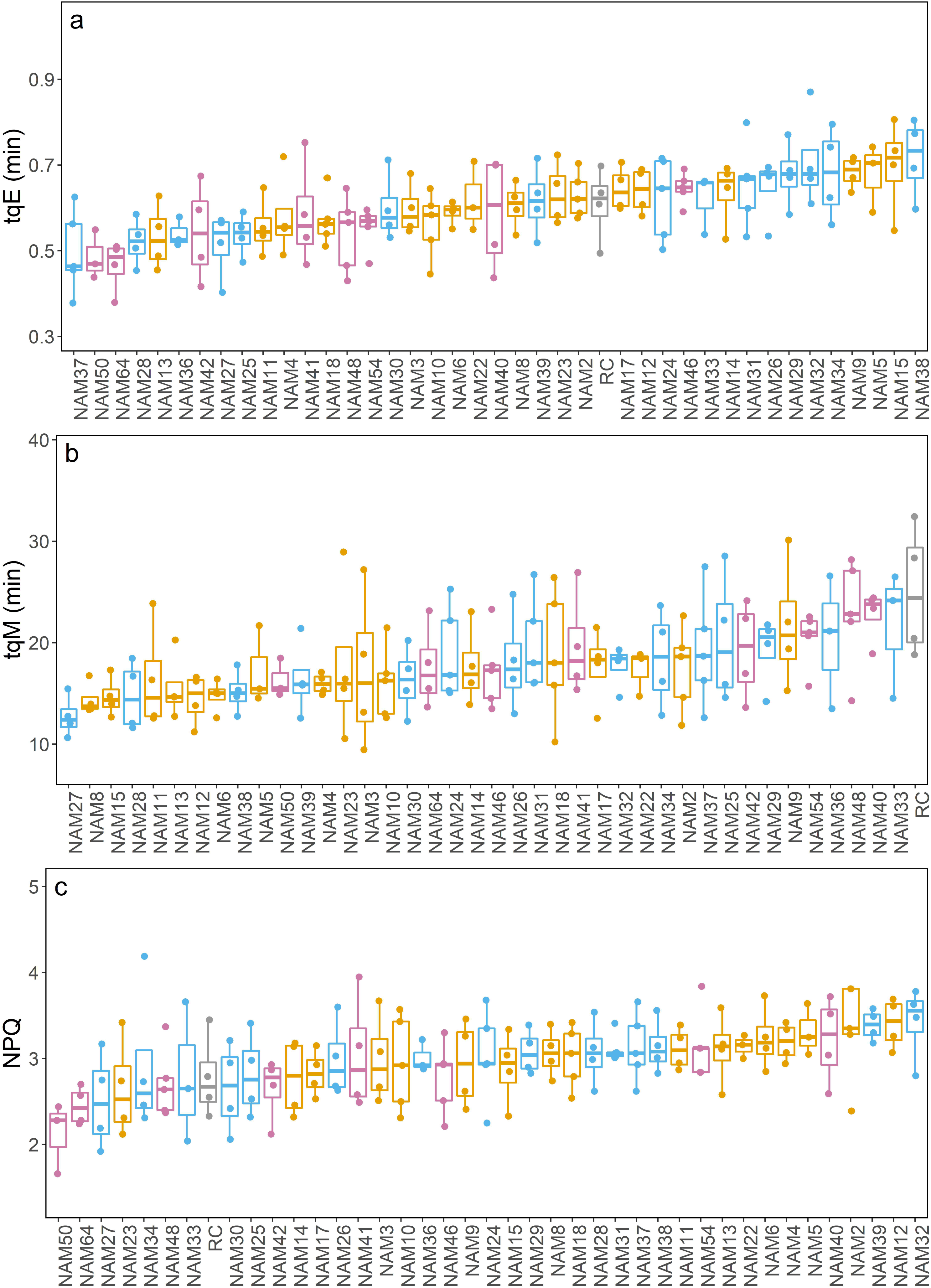
NPQ relaxation kinetics of the 41 parents of the soybean NAM population in 2019. Box plots representing (a) τqE, (b) τqM, (c) maximum NPQ, displaying mean and standard deviation. Data points for the mean of n=3-5 (See Supplemental Data 1 for details) biological replicates are displayed. Colors correspond to genotype groups, including: Reference genotype (grey), EL = elite genotype (orange), BX = breeding genotype with exotic ancestry (blue), PI = plant introductions (magenta).

Here we describe in detail the results for 2019, which was performed with increased technical replicates to allow for discarding of outliers facilitating statistical analysis, but population level trends were similar in 2018 (Supplemental Data 1; Supplemental Figure 1 and 2). In 2019, there was a statistically significant difference in the amplitude of qE (AqE) between parents (F(40,127) = 2.06, p=0.0013), which ranged between 1.57 ± 0.11 (NAM64) and 2.36 ± 0.28 (NAM2)(Figure 1). Measured NPQ attributable to qI, AqI (F(40,127) = 1.54, p=0.037), differed ~2 fold between the smallest (0.09 ± 0.05; NAM33) and largest (0.021± 0.03; NAM28) (Figure 1). In terms of relaxation rates, there was a globally statistically significant difference (F(40,127) = 2.51, p=<0.001) in measured τqE, which varied ~1.5 fold between the fastest (28.2 ± 3.6s; NAM64) and slowest (43.2 ± 5.4s; NAM38) parent, but non-significant pairwise comparisons by Games-Howells posthoc test, which may be a result of the relatively low replication. In addition, there was a ~2 fold difference in τqM between the fastest (12.7 ± 2.0 min; NAM27) and slowest relaxing (25.0 ± 6.5 min; RC) parents which was not statistically significant (p>0.05) (Figure 2; Supplemental Data 2) and there was no statistically significance difference between maximum recorded NPQ values (lowest, 2.13 ± 0.41, NAM50; highest, 3.42 ± 0.43)(Figure 2). However, taken together, these data indicate there is substantial variation in NPQ parameters within soybean germplasm.

Comparing trends from both years, there was a significant positive correlation between maximum NPQ and AqE (0.75, p<0.001 in 2018; 0.67, p<0.001 in 2019) (Supplemental Figures 3 and 4). However, there was no correlation between maximum NPQ and AqI (0.08, p>0.05 in 2018; 0.26, p>0.05 in 2019)(Supplemental Figures 3 and 4), which would be in accordance with the majority of qI being unrelated to photoprotection, being instead generated by photodamage, given the assumption NPQ increases in response to a greater requirement to dissipate excess reducing power.

While measured parameters fell within in a similar range in both 2018 and 2019 (Figure 1 and 2; Supplemental Figure 1 and 2), parents behaved differently between years, as there was no correlation between the recorded values in 2018 and 2019 (Supplemental Figures 5-10), which is consistent with a strong environmental and genotype by environment effect on NPQ. To investigate this further, we analyzed the variation in climatic conditions over the growing period in 2018 and 2019 taking data from a weather station on a nearby farm (Figure 3). Fluorescence measurements were made on day 201 in 2018 and day 207 in 2019.

**Figure 3:**
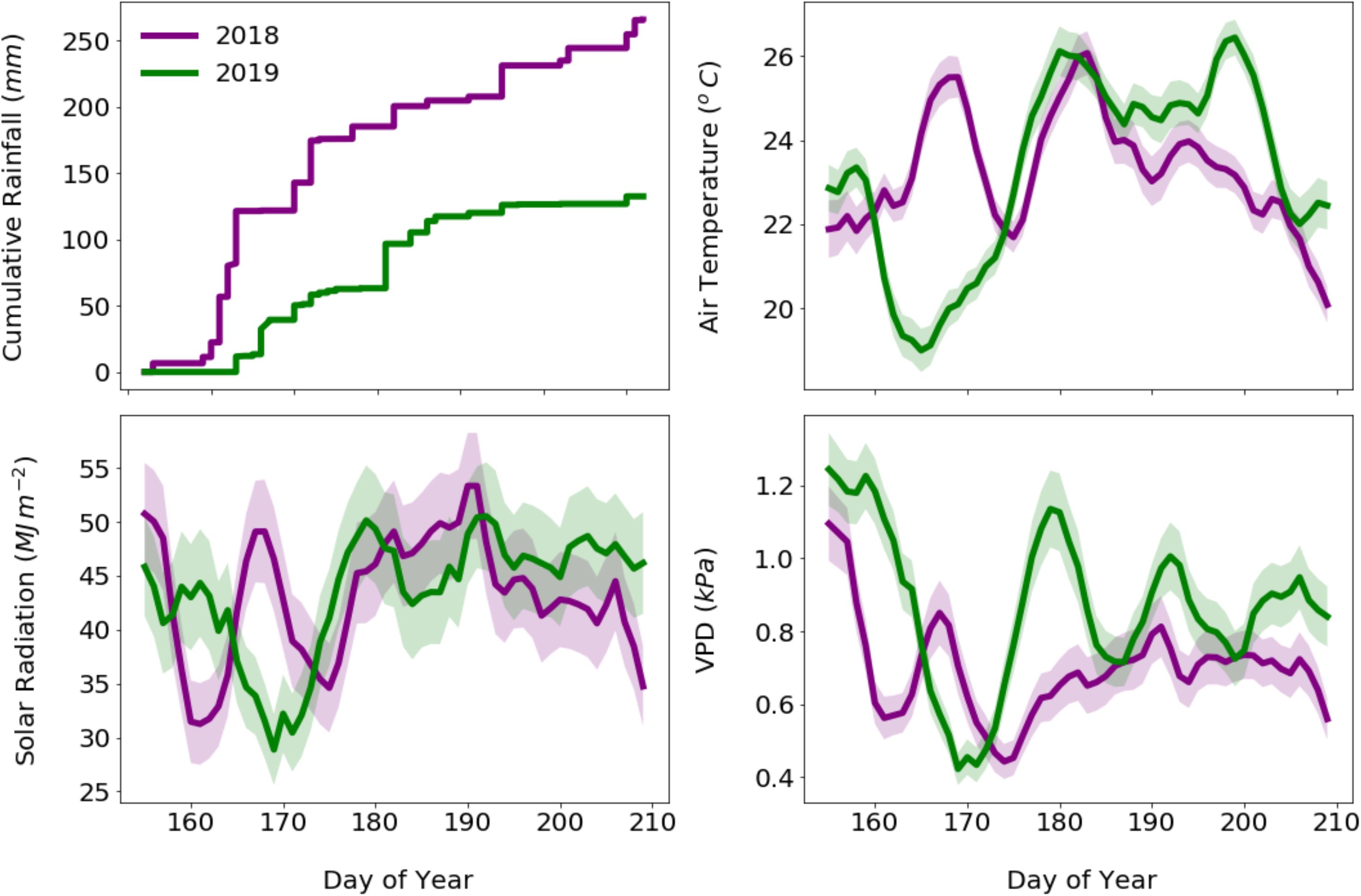
Climate data for June and July in Champaign-Urbana, Illinois comparing 2018 and 2019. Line graphs show cumulative rainfall (mm), air temperature (°C), solar radiation (MJ m^−2^) and VPD (kPa) for 2018 (purple), 2019 (green). Data are presented as 7-day running mean with +/− a 7-day running error shading.

On the day of measurement, total solar radiation and mean air temperature were similar across the two years, at ~45 MJ m^−2^ and ~23 °C respectively, while VPD was at ~0.8 in 2019 compared to ~0.7 (Figure 3). However, cumulative rainfall in 2018 was approximately double that of 2019 reaching 250 mm over June and July, and average air temperature was higher in the early growing season, increasing from 22 °C to a high of 26 °C over the first 20 days, compared to a decrease in temperature from 23 °C to 19 °C in 2019 (Figure 3). While there was no clear difference in total incident solar radiation, which is influenced by cloud cover and fluctuated between 30 and 55 MJ m^−2^ in both years (Figure 3), the differences in precipitation and temperature were reflected in periodically higher vapor pressure deficit (VPD) in 2019, which fluctuated between 1.2 and 0.4 kPa (Figure 3)

In order to assess the potential for breeding soybean for improved NPQ characteristics we calculated the heritability of parameters showing significant variation between genotypes including AqE (0.21) and AqI (0.11) using data from 2019 (Table 1).

**Table 1:**
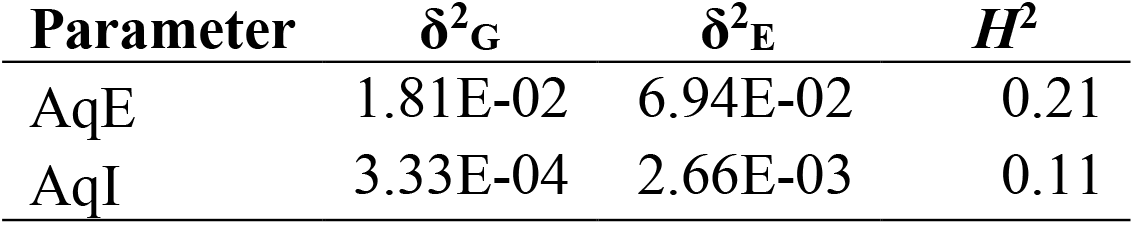
Estimates of heritability of NPQ relaxation kinetics AqE, AqI, and τqE based on field data. Data are shown for 2019, including the genotypic (δ^2^G) and environmental (δ^2^_G_) variance and the derived broad sense heritability value (*H*^2^) for AqE and AqI. Heritability values are represented to two significant figures.

## Discussion

Increasing the rate of NPQ activation and relaxation has been demonstrated as a means to improve photosynthetic efficiency (Kromdijk et al., 2016), and although further research is required to understand potential species-specific differences (Garcia-Molina and Leister, 2020) and the potential effects of reduced ROS production from quicker activation (Davis et al., 2017), targeting NPQ is proposed as a means for crop improvement (Long et al., 2019). The data collected here indicate there is a statistically significant difference in NPQ relaxation rates in the parents of the soybean NAM population, and this study provides the basis for further research to map genomic loci responsible through screening of the wider population (Song et al., 2017; Diers et al., 2018).

Taking the slowest, fastest and average short term relaxation rates of the NAM parents in 2019, and inputting them into a soybean canopy photosynthesis model suggested the difference between the slowest and average could result in a small but substantial (1.3 %) increase in carbon assimilation on a cloudy day (Wang et al., 2020). However, it remained unclear to what extent this variation in soybean is heritable, information which is required to estimate the ease with which variation in NPQ could be selected for in a conventional breeding program. We therefore sort to address this through analysis of the entire NAM population.

NPQ is a multigene trait, and is likely to be affected by environmental fluctuations that influence cellular energy status, including temperature, water availability, nutrient availability and pathogen attack. Subsequently, we observed a low (< 0.5) heritability value for NPQ relaxation parameters in the NAM population using current screening approaches in a field environment. This is similar to heritability estimates from OJIP measurements of chlorophyll fluorescence in a larger panel of soybean genotypes (Herritt et al., 2018), and those reported for other photosynthetic traits such as *J*max in wheat (0.32; (Driever et al., 2014)).

In addition, we observed no significant correlation between measured parameters over 2018 and 2019, suggesting a strong environmental effect. Meteorological data indicates conditions on the day of sampling were similar, but plants may have been more water limited throughout the duration of the growing season in 2019. Therefore, the environmental cause of variation in NPQ relaxation parameters between 2018 and 2019 could be related to long term adaptation to annual differences in rainfall and temperature, requiring further investigation. However, heritability is also dependent on the presence of large effect alleles within a population. While significant genotypic variation in AqE and AqI was identified in the NAM population, the data also suggest that unlike in some other species, such as rice (Kasajima et al., 2011), there were no large effect alleles present in the population screened.

Therefore, several approaches could be considered to improve the potential for breeding faster relaxation of NPQ: screening a bigger collection of diverse soybean genotypes to identify potential large effect alleles, growing and measuring genotypes under controlled conditions to reduce environmental variation, increasing replication over multiple environments, and improving statistical methods to reduce variation. One step towards achieving this goal could be coupling large scale screening with open availability of data as reported elsewhere (Kuhlgert et al., 2016).

Genetic modification provides an alternative means of modulating NPQ (Armbruster et al., 2016; Kromdijk et al., 2016), and to be successful this will depend on both an understanding of the genes regulating NPQ relaxation, and how they influence plant energetics in a fluctuating environment (Kramer and Evans, 2011). While several of the genes involved in fast and medium term relaxation of NPQ have been identified, further work is required to understand which are the key genes and processes underlying long term NPQ relaxation, or qI. Advances in what in addition to new forward genetic screens, such as those that identified the novel slow relaxing phase qH (Malnoë et al., 2018), and refinement of candidate gene lists identified by genome wide association analysis (Rungrat et al., 2019) is required.

In conclusion, we measured NPQ relaxation in the 41 parents of the soybean NAM population; while there is significant genotypic variation, there was a strong environmental effect resulting in low heritability estimates which imply selection by breeding will be difficult. Therefore given current approaches and information, the most effective means of manipulating NPQ for improved photosynthetic performance in soybean will likely rely on genetic modification.

## Acknowledgements

We thank Troy Cary for setting up the soybean NAM population plots, Wanne Kromdijk for help setting up scripts for analysis of NPQ relaxation parameters and Liana Acevedo-Siaca for support with statistics. This work was supported by the project Realizing Increased Photosynthetic Efficiency (RIPE), funded by the Bill and Melinda Gates Foundation (BMGF), Foundation for Food and Agriculture Research (FFAR) and the UK Department for International Development (UKAid) under grant number OPP117215. SJB was supported by Carl R. Woese Institute for Genomic Biology Fellowship and EMDB is supported by an Illinois Distinguished Fellowship.

## Author contributions

SPL, SJB and BWD designed the study. SJB, EMBD, IC, IF and SC conducted the NPQ measurements, CM conducted the meteorological data collection. SJB and CM performed the data analysis. SJB, SPL, EMDB, CM and BWD wrote the article and all authors provided critical revisions.

