## Supplementary figures and images for "Variation in relaxation of non-photochemical quenching in a soybean nested association mapping panel as a potential source for breeding improved photosynthesis"

### Supplemental Figure 3

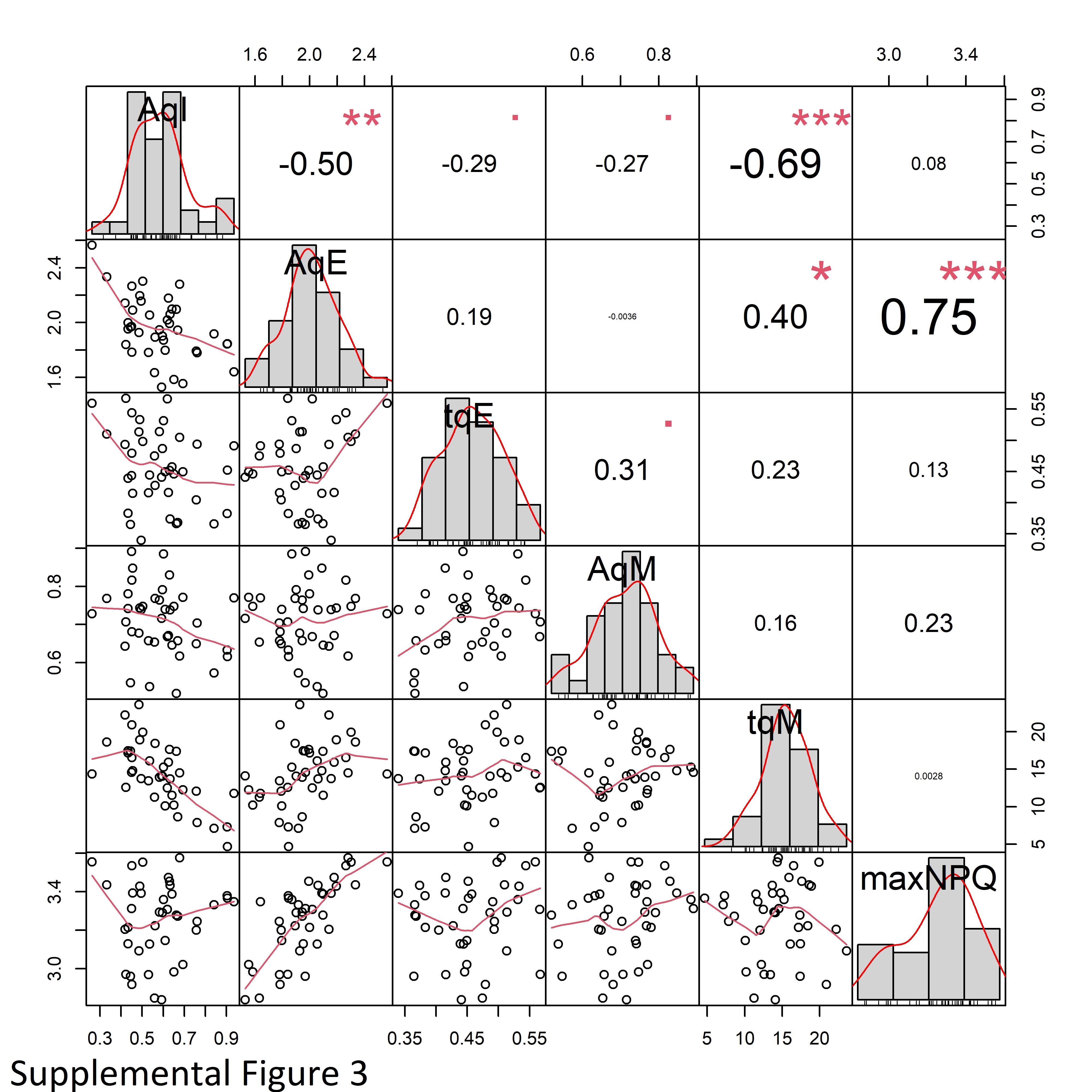

### Supplemental Figure 4

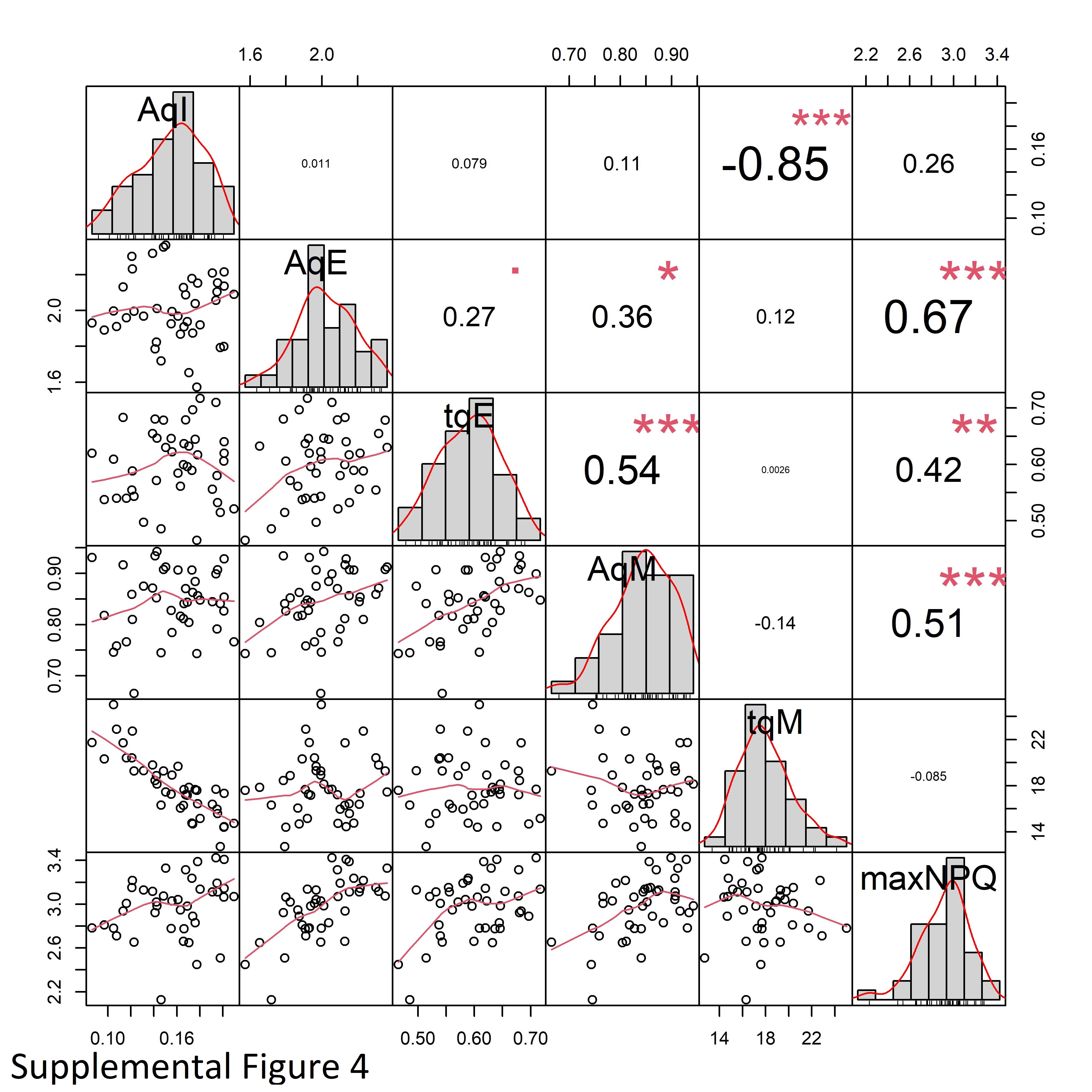

### Supplemental Figure 5

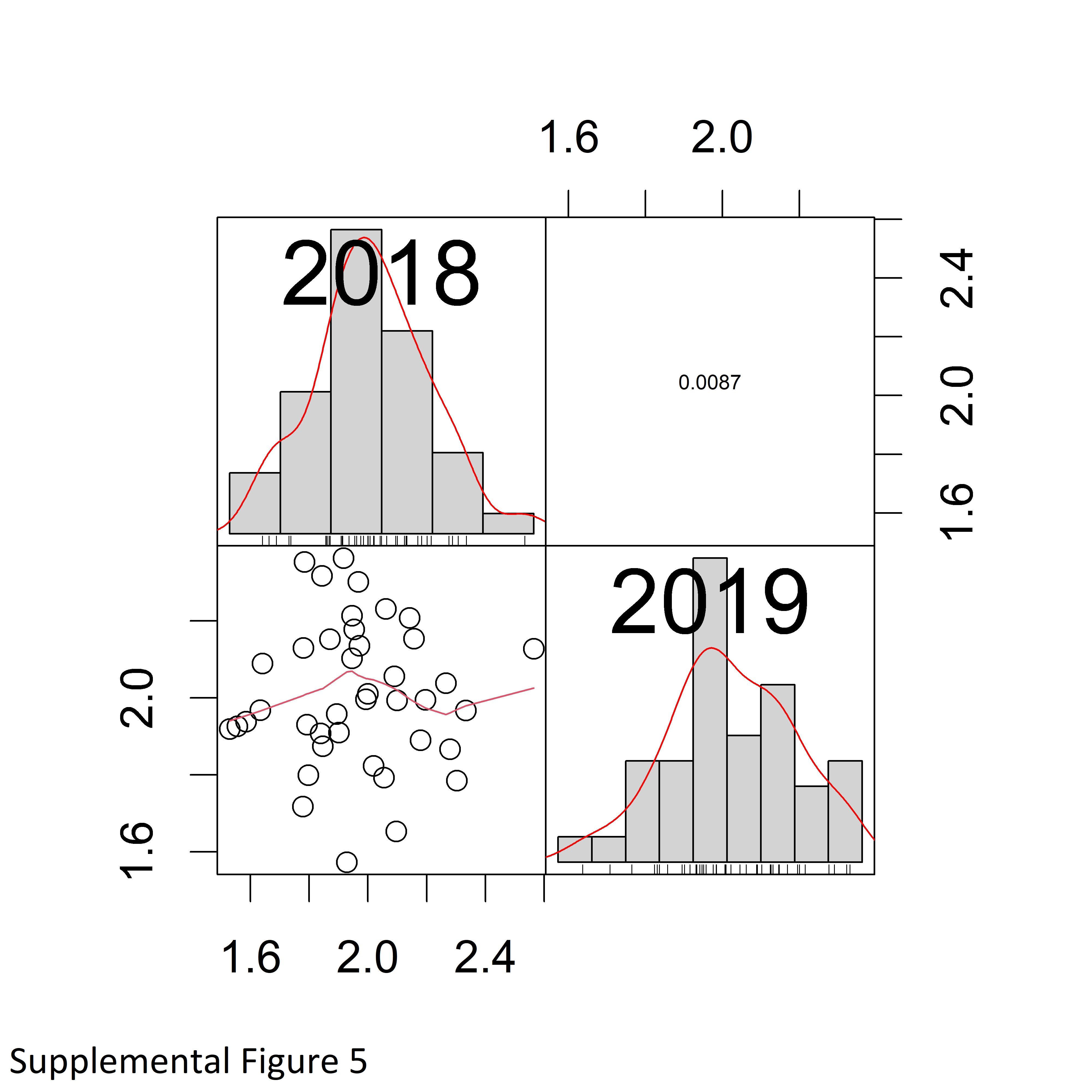

### Supplemental Figure 6

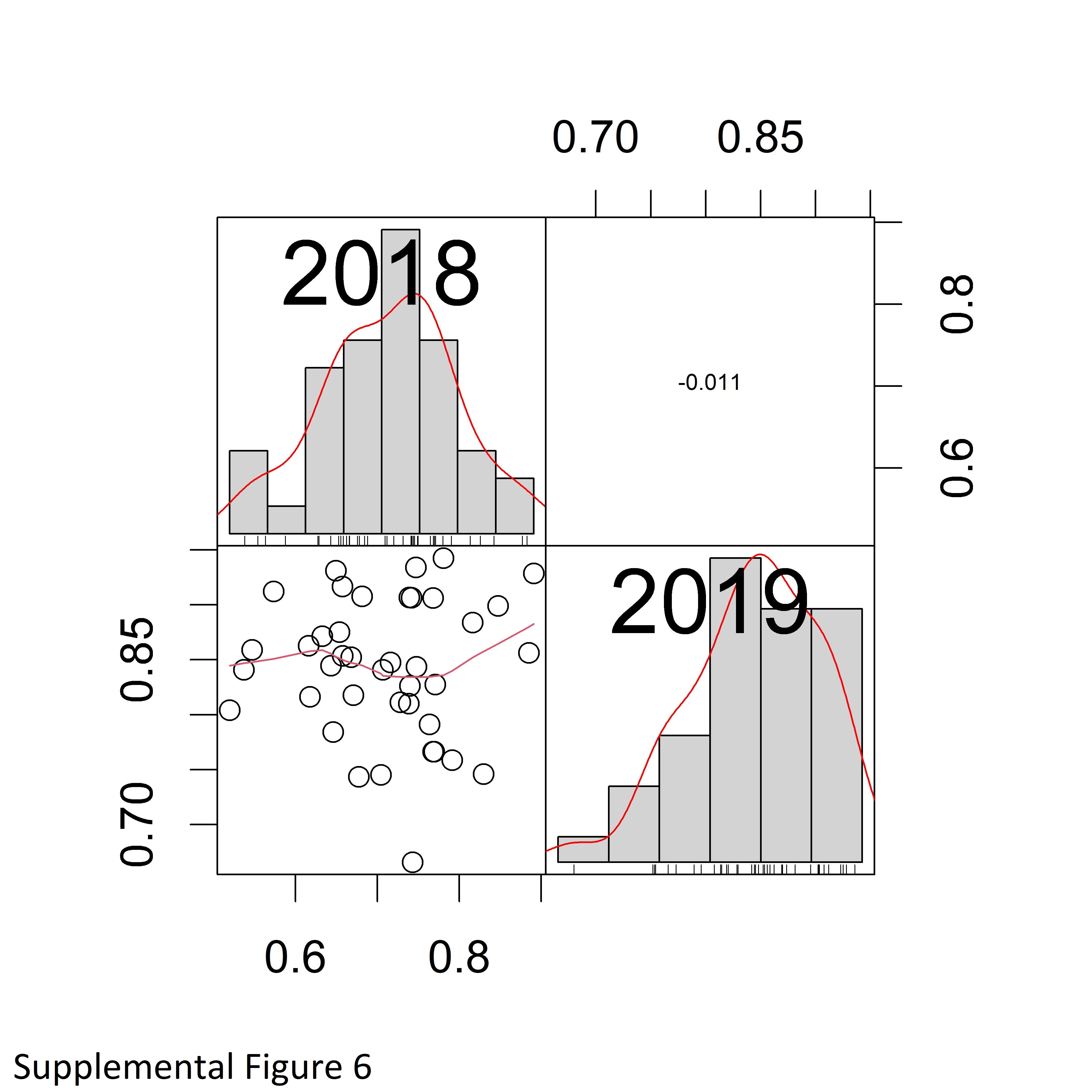

### Supplemental Figure 7

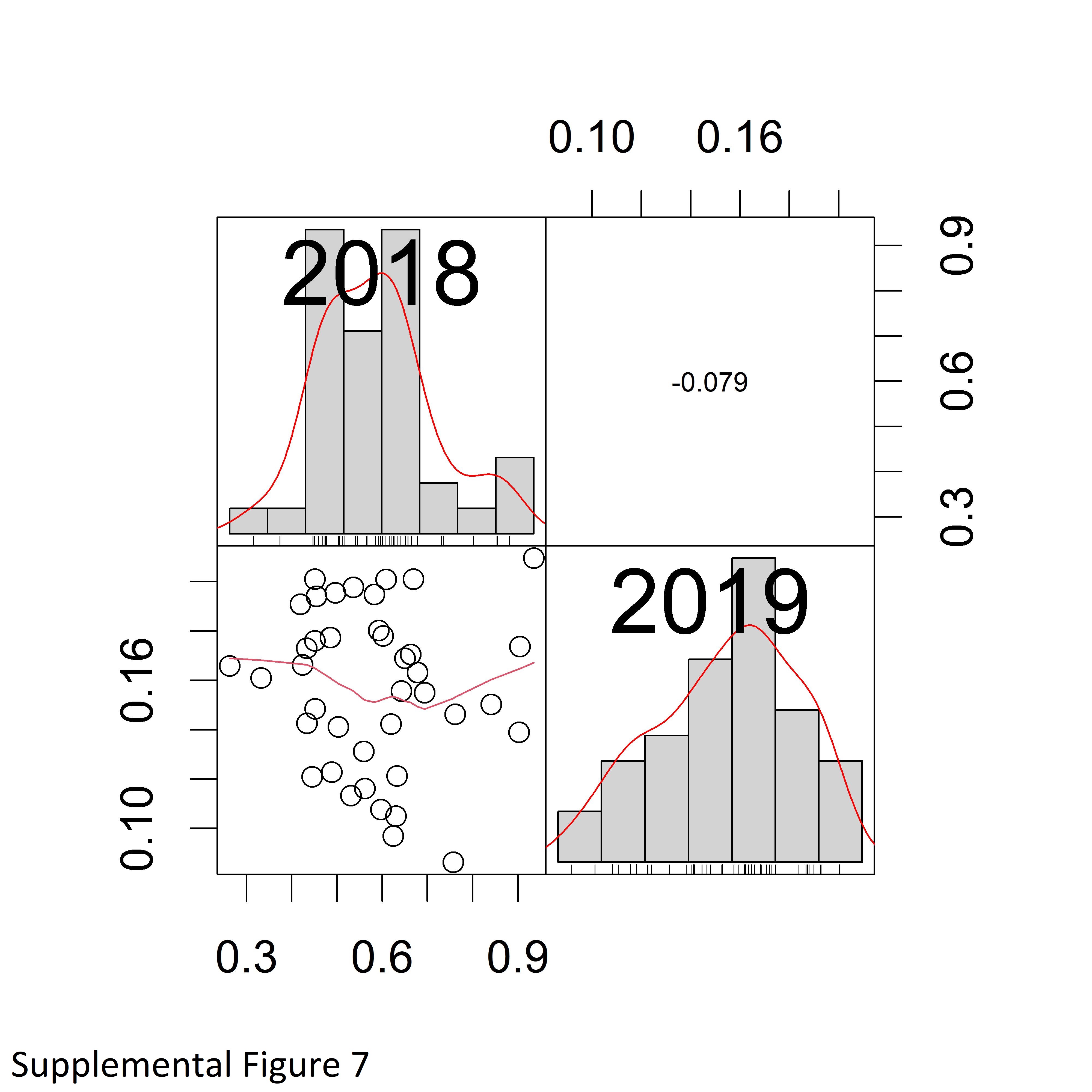

### Supplemental Figure 8

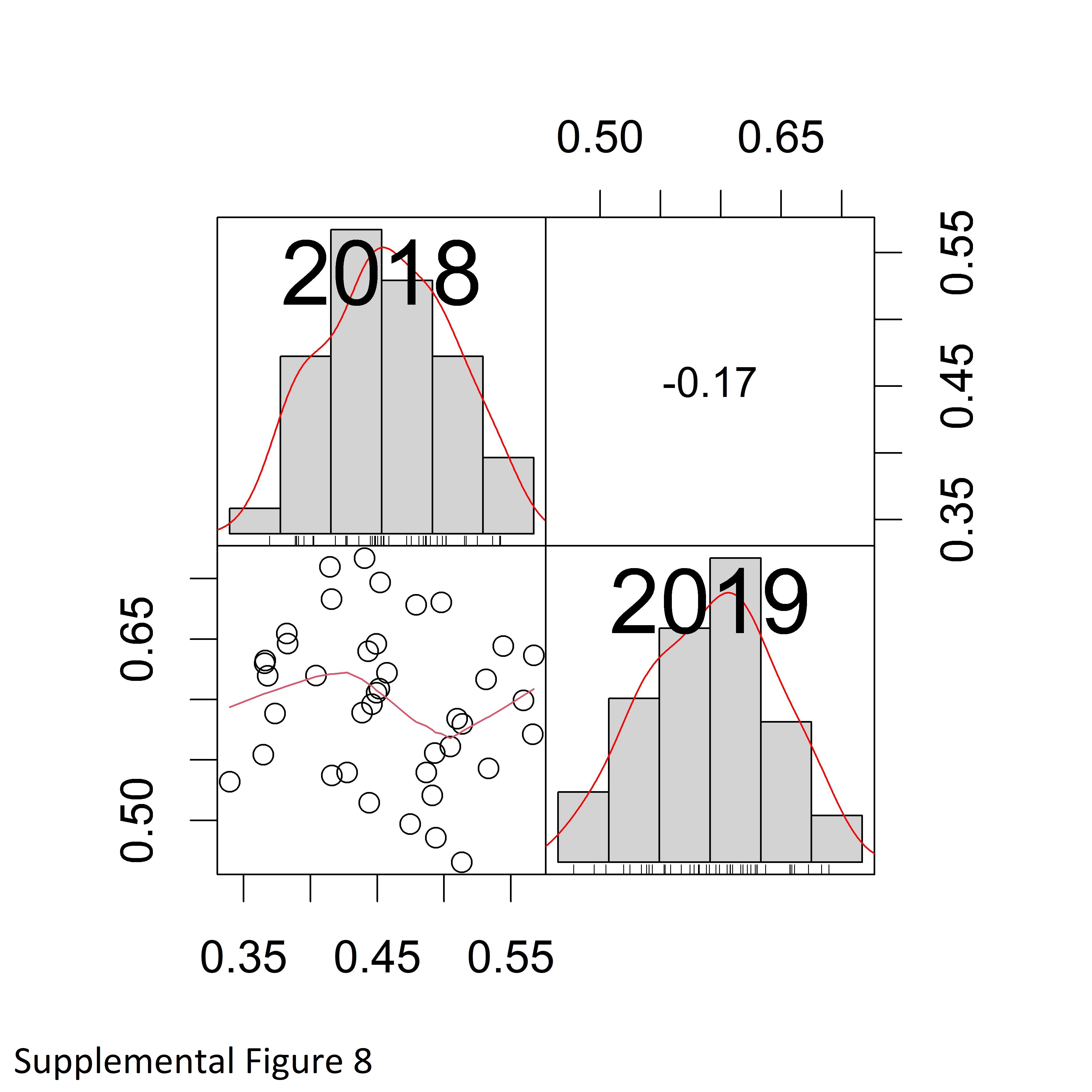

### Supplemental Figure 9

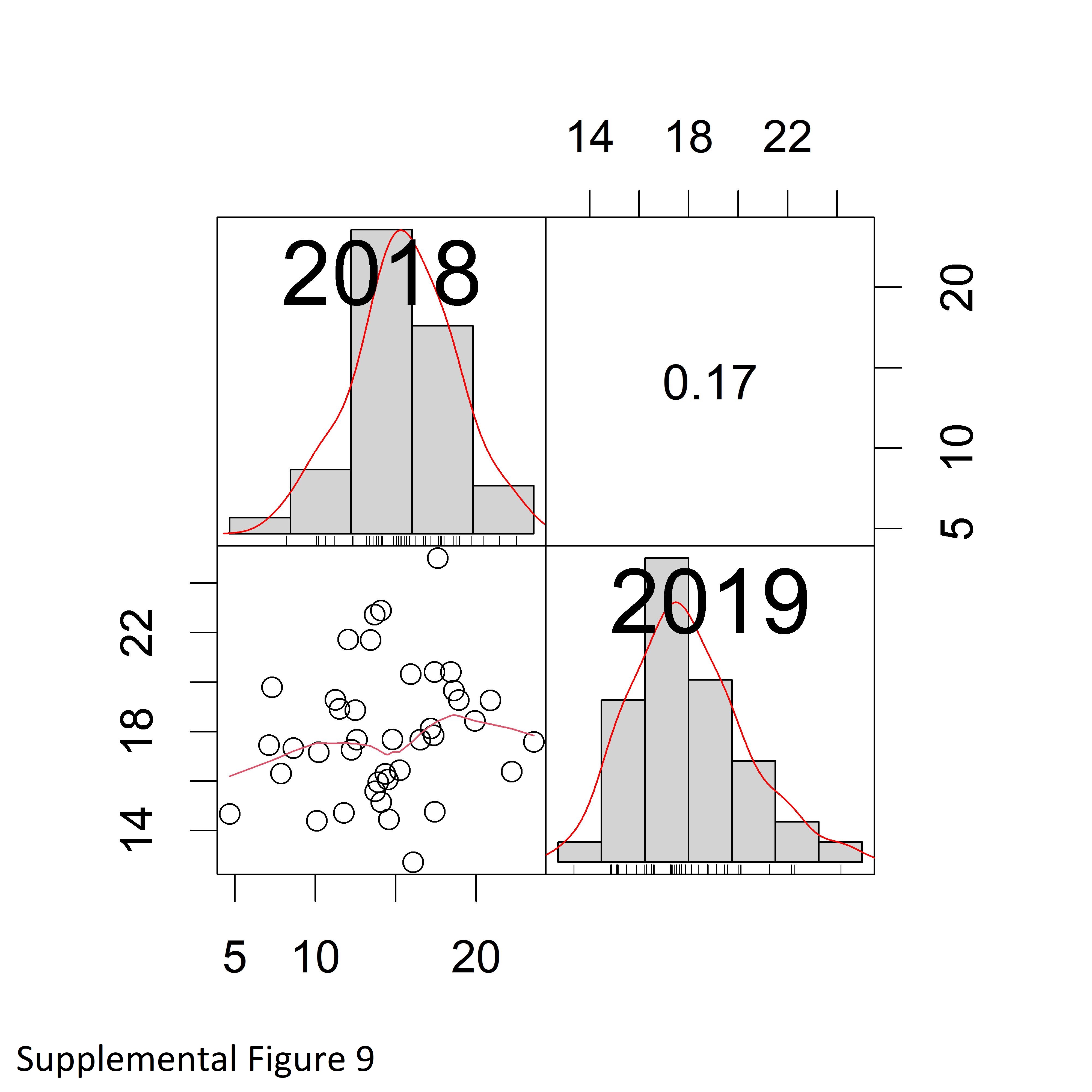

### Supplemental Figure 10

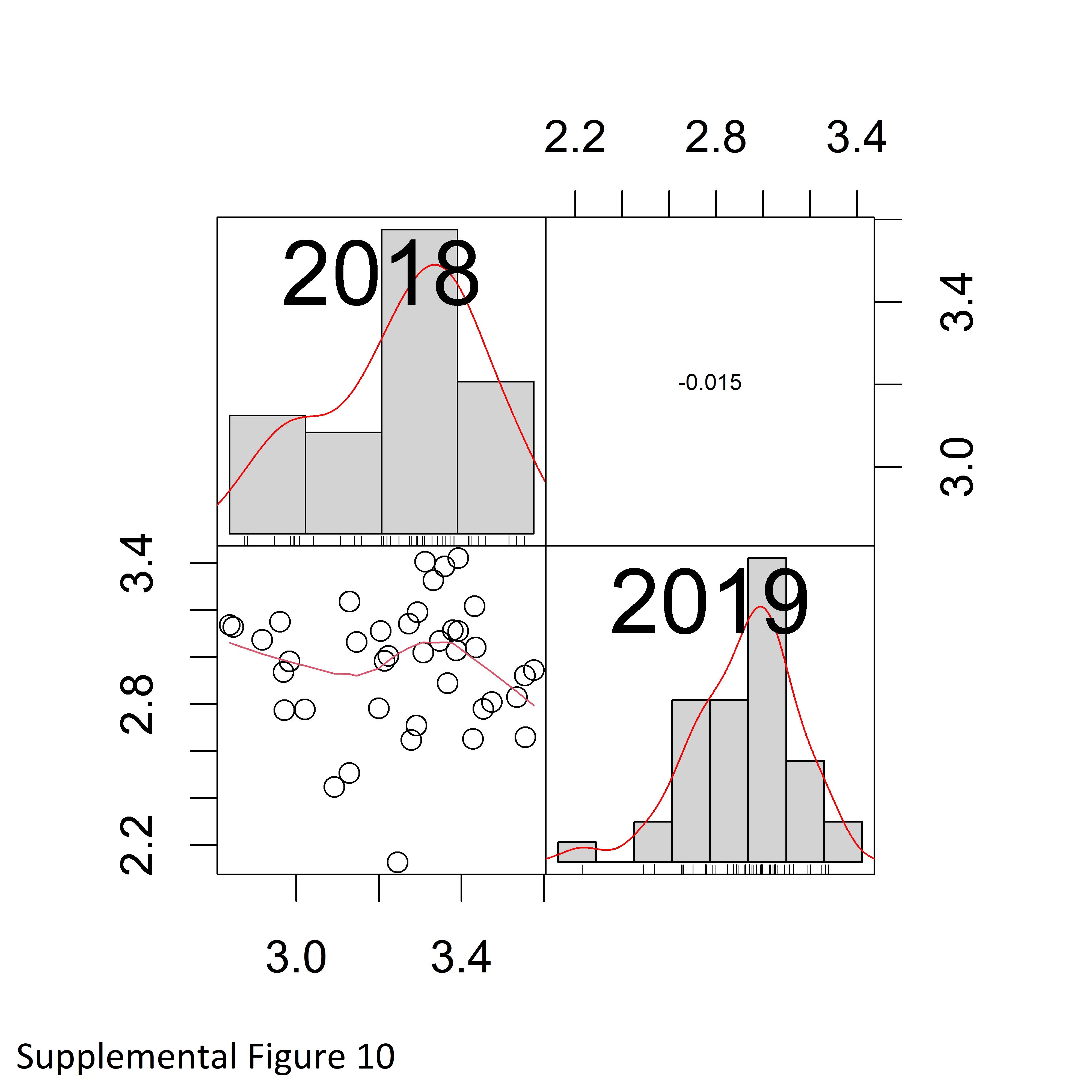

### Supplemental Figure - 1

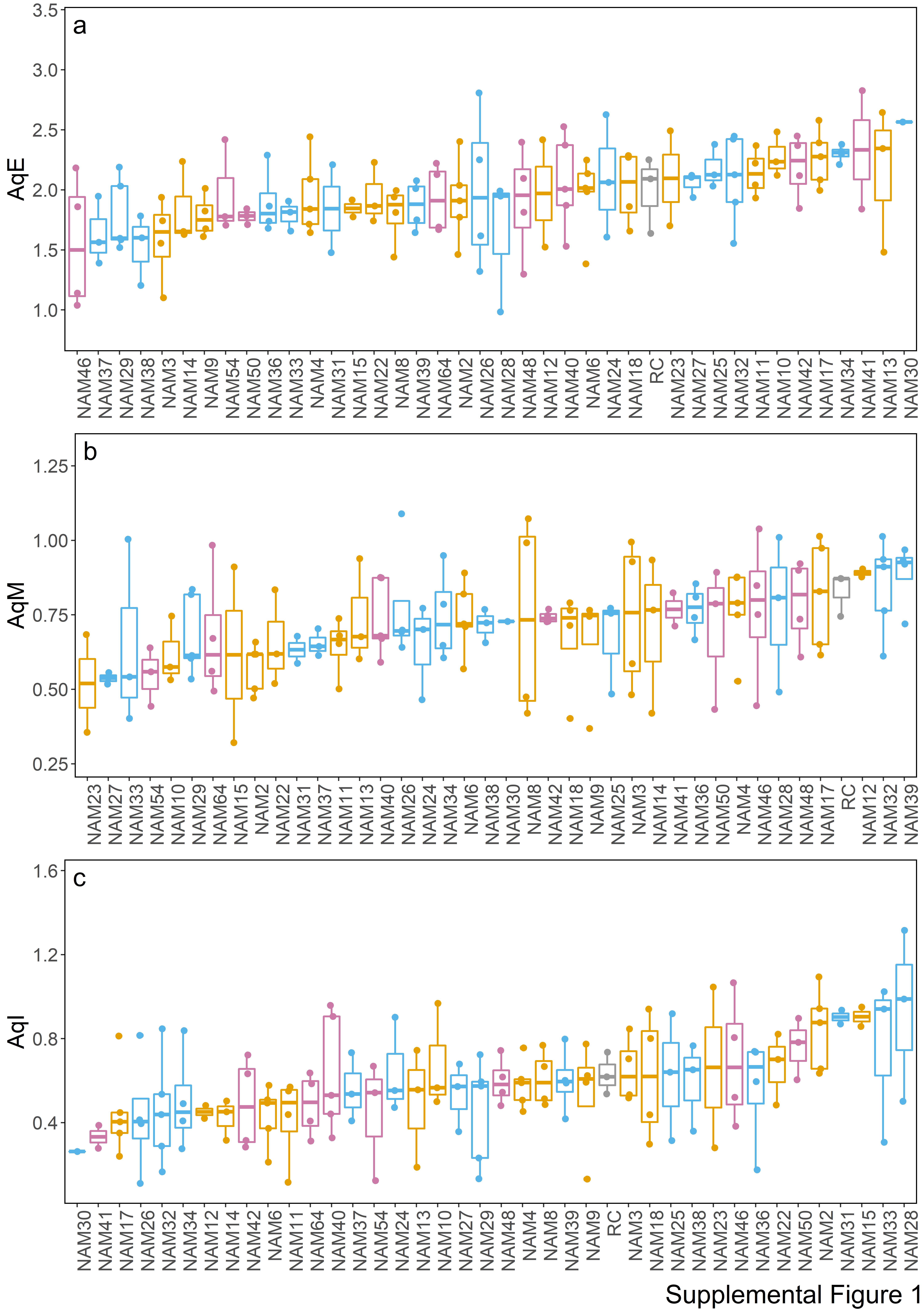

### Supplemental Figure - 2

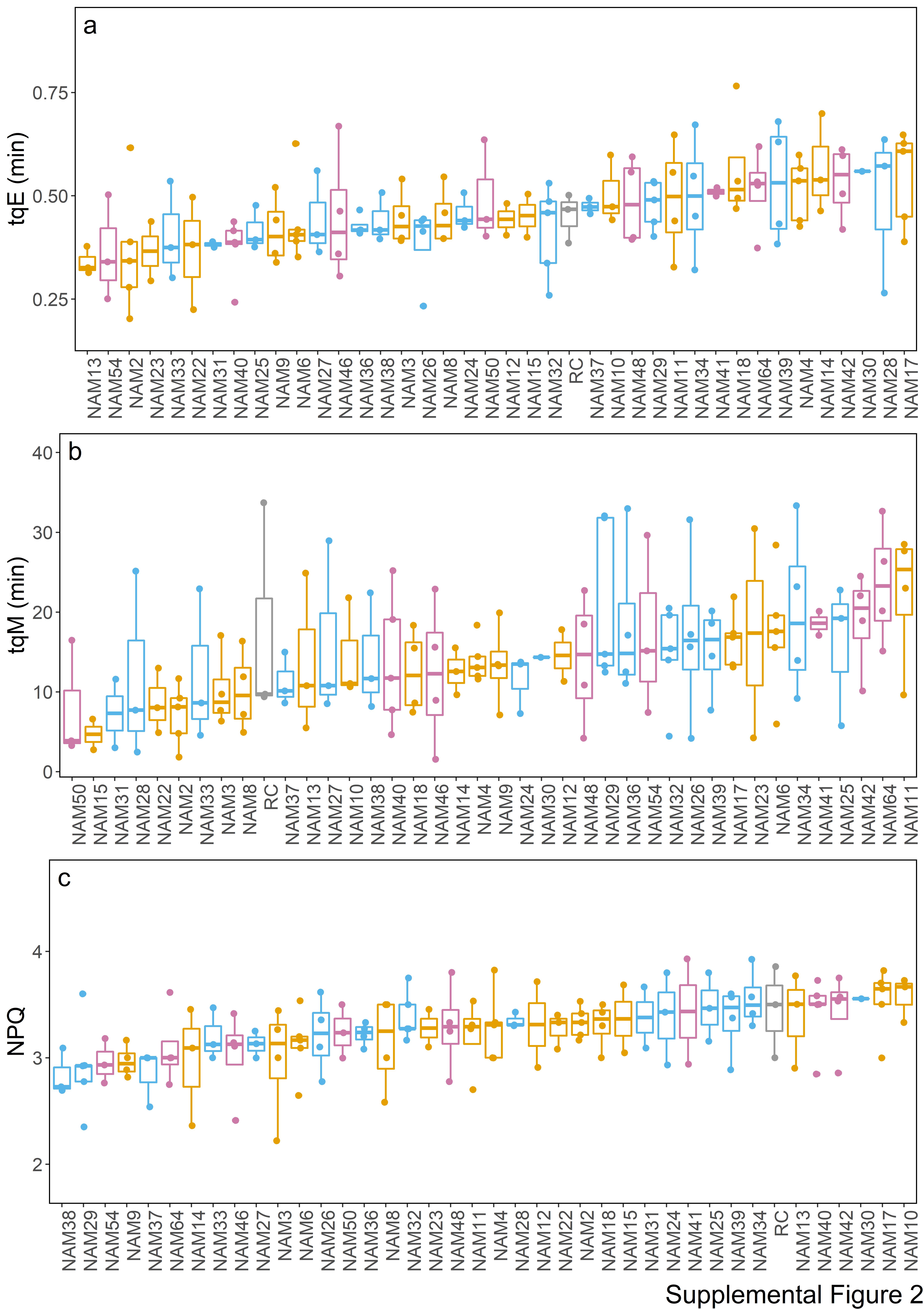
